# Highly redundant neuropeptide volume co-transmission underlying episodic activation of the GnRH neuron dendron

**DOI:** 10.1101/2020.08.25.266080

**Authors:** Xinhuai Liu, Shel-Hwa Yeo, H. James McQuillan, Michel K. Herde, Sabine Hessler, Isaiah Cheong, Robert Porteous, Allan E. Herbison

**Author notes:** Correspondence and requests for materials should be addressed to Allan E. Herbison.

## Abstract

The necessity and functional significance of neurotransmitter co-transmission remains unclear. The glutamatergic “KNDy” neurons co-express kisspeptin, neurokinin B (NKB) and dynorphin and exhibit a highly stereotyped synchronized behavior that reads out to the gonadotropin-releasing hormone (GnRH) neuron dendrons to drive episodic hormone secretion. Using expansion microscopy, we show that KNDy neurons make abundant close but non-synaptic appositions with the GnRH neuron dendron. Confocal GCaMP6 calcium imaging demonstrated that, of the neurotransmitters co-expressed by KNDy neurons, only kisspeptin was able to activate the GnRH neuron dendron. The selective deletion of kisspeptin from KNDy neurons resulted in mice in which the synchronized behavior of the KNDy neurons was maintained but their ability to drive episodic hormone secretion was abolished. This indicates that KNDy neurons drive episodic hormone secretion through converse modes of highly redundant neuropeptide co-transmission orchestrated by differential postsynaptic neuropeptide receptor expression at their two target sites.

## Introduction

Many neurons use the co-transmission of classical small molecule and neuropeptide neurotransmitters to signal within their networks. Such co-transmission enables a wide dynamic range of signaling through the frequency coding of transmitter release (van den Pol 2012, Vaaga, Borisovska et al. 2014, Tritsch, Granger et al. 2016). However, the extent and functional significance of co-transmission remains unclear in most forebrain circuits. For example, “Dale’s Principle”, as formulated by Eccles (Eccles 1976), posits that all axons of an individual neuron will release the same set of transmitters. The generality of this concept has now been challenged for small molecule co-transmitters (see (Tritsch, Granger et al. 2016)) but remains untested for neuropeptide co-transmission.

The kisspeptin neurons located in the arcuate/infundibular nucleus of the mammalian hypothalamus appear to engage in substantial co-transmission being glutamatergic and synthesizing at least four neuropeptides including kisspeptin, neurokinin B, dynorphin and galanin (Lehman, Coolen et al. 2010, Skrapits, Borsay et al. 2015). In a range of experimental mammals, the majority of these arcuate nucleus (ARN) neurons co-express Kisspeptin, Neurokinin B and Dynorphin resulting in their “KNDy” moniker (Lehman, Coolen et al. 2010, Skrapits, Borsay et al. 2015). Immunohistochemical studies have demonstrated that the three neuropeptides are packaged within separate vesicles within KNDy nerve terminals (Lehman, Coolen et al. 2010, Murakawa, Iwata et al. 2016). Although the KNDy neurons project widely throughout the limbic system (Krajewski, Burke et al. 2010, Yeo and Herbison 2011, Yip, Boehm et al. 2015), they are best characterized as being the “GnRH pulse generator” responsible for episodically activating the gonadotropin-releasing hormone (GnRH) neurons to drive pulsatile luteinizing hormone (LH) secretion (Clarkson, Han et al. 2017, Herbison 2018, Plant 2019). The KNDy neurons are proposed to achieve this by providing an episodic stimulatory input to the distal projections of the GnRH neurons close to their secretory zone in the median eminence (Herbison 2018). These distal projections of the GnRH neuron have shared features of dendrites and axons and have been termed “dendrons” (Herde, Iremonger et al. 2013, Herbison 2016, Moore, Prescott et al. 2018). The nature and functional significance of KNDy co-transmission at the GnRH neuron dendron remains unknown.

## Results

### KNDy neurons form abundant close appositions with GnRH neuron distal dendrons

We first established the anatomical relationship between KNDy fibers and the GnRH neuron distal dendrons in the ventrolateral ARN using confocal immunofluorescence. Analysis of horizontal sections (Suppl.Fig.1) revealed numerous kisspeptin-expressing fibers passing through and around the GnRH neuron dendrons as they turned towards the median eminence (Fig.1A). We found that 68.5±6.2% of analyzed GnRH dendrons segments had at least one apposition with a kisspeptin fiber (N=4 female mice). To examine fibers originating from KNDy neurons, we assessed the relationship of GnRH neuron dendrons to fibers co-expressing kisspeptin and NKB. In total, we observed 4.0±0.4 kisspeptin close appositions/100μm of dendron length and 67.0±7.6% of these co-expressed NKB (Fig.1A; N=4). Prior work has shown that the GnRH neuron projections are innervated by KNDy neurons as well as preoptic area kisspeptin neurons that do not express NKB (Yip, Boehm et al. 2015). On average, each kisspeptin/NKB fiber made close appositions with 4.5±0.7 GnRH neuron dendrons. This indicates an abundant, convergent innervation of the GnRH neuron distal dendrons by KNDy neurons.

### KNDy neurons signal through volume transmission to the GnRH neuron dendrons

While regular confocal analysis is useful for assessing anatomical relationships, it is unable to unambiguously define synapses. Expansion microscopy (ExM) uses isotropic swelling of the tissue specimen to provide ∼70 nm spatial resolution that can reliably image synapses in the brain (Wassie, Zhao et al. 2019). We have previously demonstrated using ExM that a “side-plane” overlap of >0.23 μm (0.95 μm post-expansion) or face-plane (z stack) overlap >0.42 μm (1.75 μm post-expansion) between a synaptophysin-immunoreactive bouton and the GFP of a GnRH neuron represents a *bona fide* synapse (Wang, Guo et al. 2020). The presence of kisspeptin synapses on distal dendrons was examined by assessing synaptophysin-containing kisspeptin boutons opposed to GFP-expressing dendrons. Surprisingly, we identified no kisspeptin-containing synaptic profiles on 45 individual distal dendrons (> 15 µm length each, N=3 mice) with all kisspeptin/synaptophysin-immunoreactive boutons being outside the criteria for a synapse or indeed quite separate from the GFP-expressing dendron (Fig.1B and 1C). The average distance of apparently opposing kisspeptin-synaptophysin boutons from GnRH neuron dendrons was 2.22±0.27 µm (post-expansion). Nevertheless, many synaptophysin-expressing boutons without kisspeptin were identified to make synaptic appositions with GnRH neuron dendrons (density of 2.3±0.1 synaptophysin synapses per 10µm dendron)(Fig1B). This suggested that kisspeptin inputs to the distal dendron did not make conventional synapses. To verify this, we used the same approach to examine the morphological relationship of kisspeptin inputs to the GnRH neuron soma/proximal dendrites in the rostral preoptic area, where local kisspeptin neurons form conventional synapses with GnRH neurons (Piet, Kalil et al. 2018). We assessed eight 60 μm-lengths of GFP soma/dendrite in each of three mice and found many synaptophysin-kisspeptin boutons forming synapses with GFP-expressing dendrites (Fig.1D). Overall, 37.9 ± 0.8% of all synaptophysin boutons synapsing on GnRH neuron soma/proximal dendrites (0.81±0.44 per 10 μm) contained kisspeptin.

These observations indicate that while kisspeptin inputs to the GnRH cell bodies and proximal dendrites exist as classical synapses, this is not the case for the distal dendron where KNDy signaling occurs through short-diffusion volume transmission.

### GnRH neuron dendrons only respond to one of the four KNDy co-transmitters

We have previously established an acute horizontal brain slice preparation (Suppl. Fig.1A) in which changes in [Ca^2+^] within the thin GnRH neuron distal dendrons can be measured providing a proxy for electrical activity (Iremonger, Porteous et al. 2017). A thick horizontal brain slice containing the median eminence and adjacent hypothalamic tissue was prepared from adult male and diestrous-stage female *Gnrh1-Cre* mice previously given preoptic area injections of AAV9-CAG-FLEX-GCaMP6s (Suppl. Fig 1B). To mimic the episodic release of transmitters, multi-barrelled pipettes were used to apply candidate neurotransmitters as short 90s puffs to the region of the GnRH neuron dendrons while recording calcium signals from multiple dendrons simultaneously using confocal imaging. In vivo recordings show that KNDy neurons exhibit synchronized episodes of activity for 1-2 min prior to each LH pulse (Han, Kane et al. 2019, McQuillan, Han et al. 2019). In controls, where we applied artificial cerebrospinal fluid (aCSF) at different distances and pressures from the brain slice, we found that slight movement artefacts were unavoidable. Placing a pipette 30-130 μm above the surface of the brain slice and puffing aCSF resulted in a 4.5±0.5% (mean±SD) increase in [Ca^2+^] within the dendrons beneath (Fig.2A,B). As such, we defined a threshold for a drug-induced change in [Ca^2+^] as requiring an increase or decrease greater than the aCSF mean plus 2 standard deviations from the control change (i.e. >5.5%) and a response that outlasted the time of the 20s or 90s puff.

**Figure 1.**
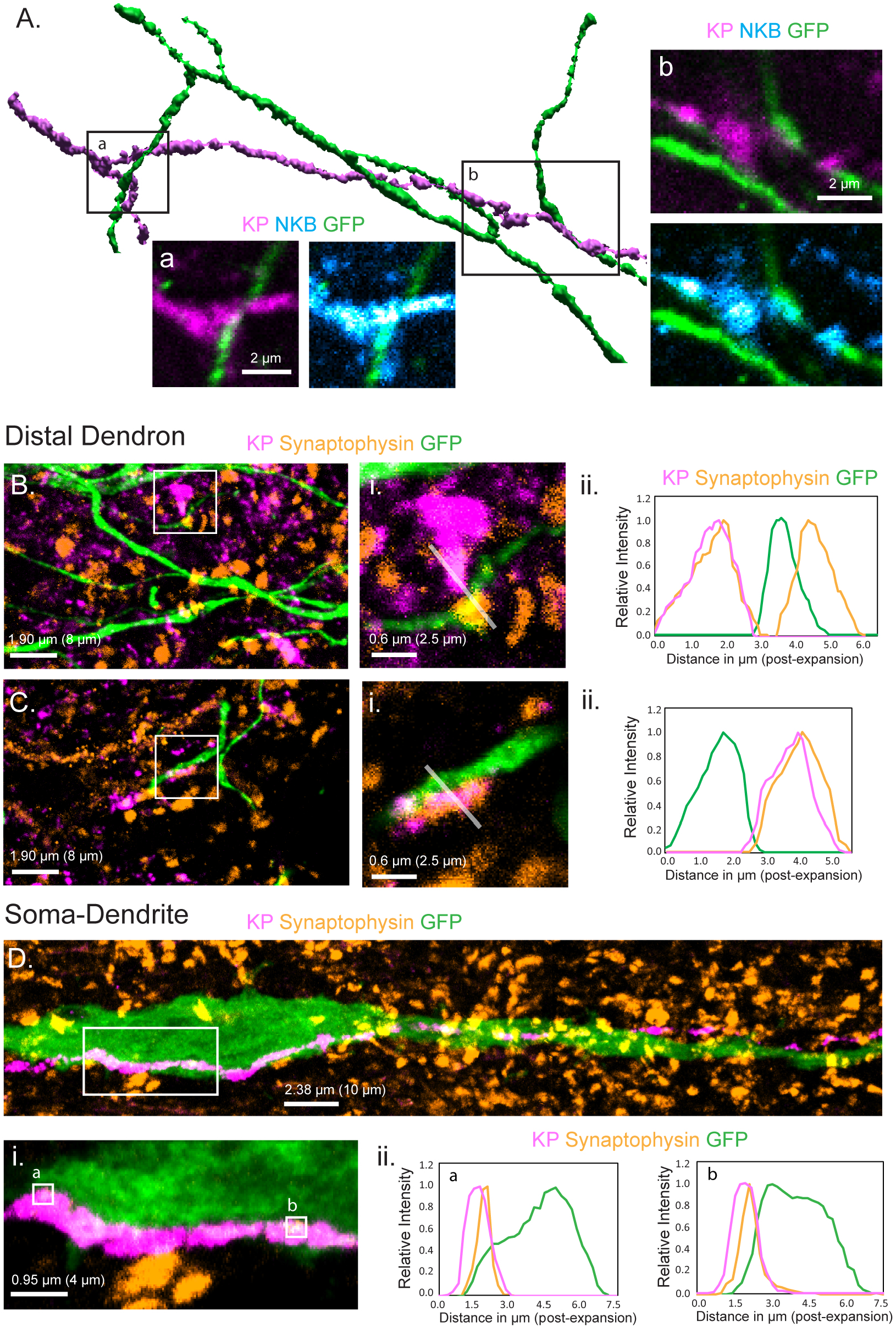
Relationship of KNDy neuron fibers to GnRH neuron distal dendrons. **A**. 3D reconstruction of regular confocal images showing a KNDy fiber expressing kisspeptin (KP) and neurokinin B (NKB) making close appositions with three GnRH neuron dendrons in the ventrolateral ARN of a GnRH-GFP mouse. **B&C**. Expansion microscopy views of GnRH distal dendrons surrounded by kisspeptin fibers and synaptophysin puncta. Insets (i.) highlight two examples of synaptophysin-expressing kisspeptin terminals adjacent to GnRH dendrons. Grey lines indicate the line scans used to generate the fluorescence relative intensity profiles shown to the right. **Bii**. shows a GFP-expressing dendron with a chemically-unidentified synapse on one side (>0.95 μm overlap between synaptophysin and GFP signals) and a kisspeptin terminal making a close non-synaptic contact on the other (no overlap). Cii. another example of kisspeptin terminal (kisspeptin and synaptophysin) making a close non-synaptic (overlap <0.95 μm) contact with a GnRH dendron. **D**. Expansion microscopy view of a GnRH neuron cell body and proximal dendrite surrounded by synaptophysin puncta and with a kisspeptin fiber running along its length. Imaging in the z-axis face view shows two locations (i.a&b) where kisspeptin/synaptophysin puncta make synapses on the GnRH neuron cell body. **Dii**. Fluorescence relative intensity profiles show two synaptophysin-containing kisspeptin boutons exhibiting >1.75 μm overlap with cytoplasmic GFP of the GnRH neuron. Scale bars show pre-expansion units with post-expansion values in brackets.

**Figure 2.**
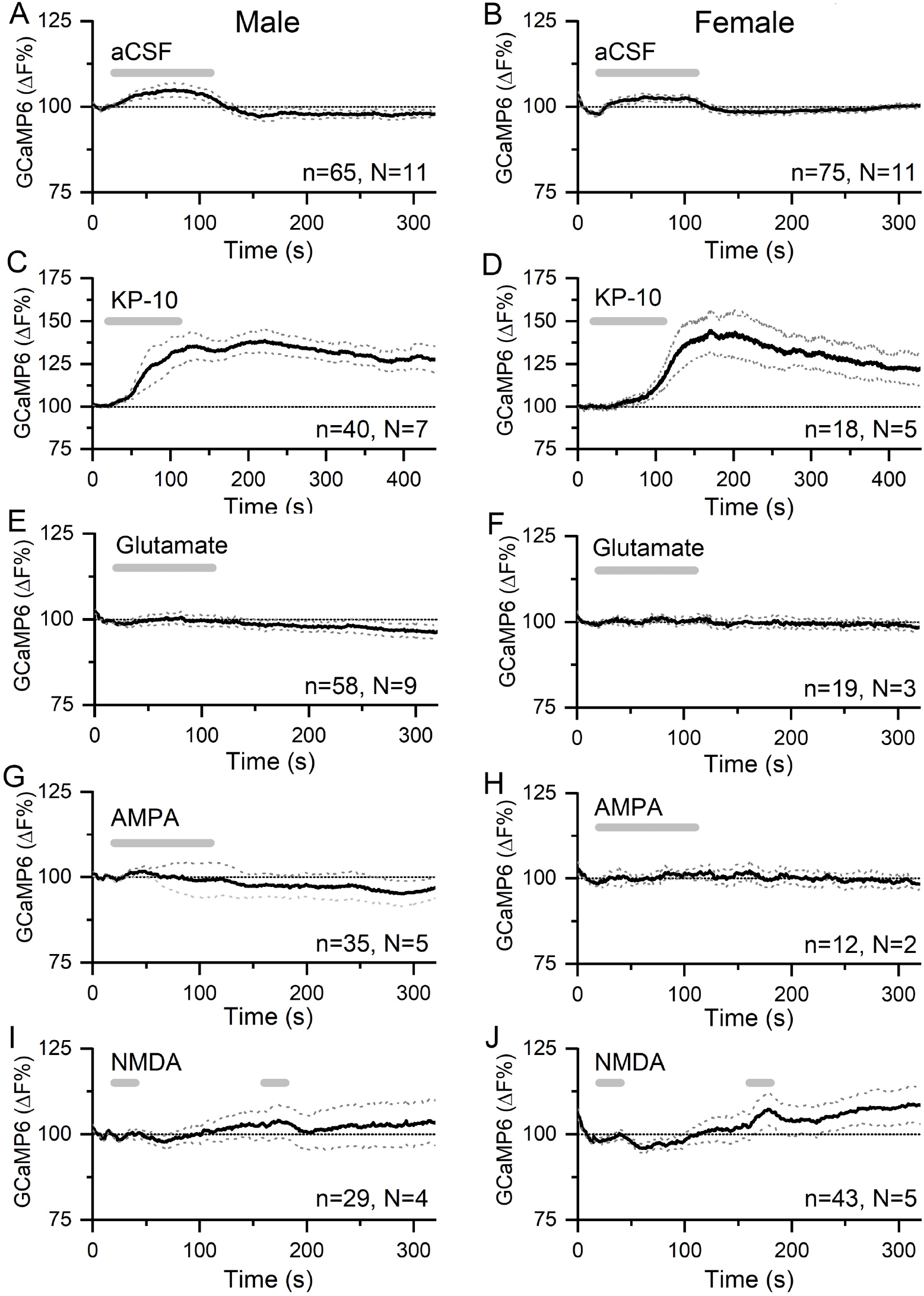
Kisspeptin but not glutamate regulates [Ca^2+^] in GnRH neuron distal dendrons. **A&B**, effects of 90s puffs of aCSF on GCaMP6 fluorescence in GnRH neuron distal dendrons in male and female *Gnrh1-Cre::GCaMP6s* mice. **C&D**, puffs of kisspeptin-10 (100nM) generate large, sustained increases in [Ca^2+^] in both sexes. Note the altered x- and y-axes. **E-J**, long (90s) or short (20s) puffs of glutamate (600μM), AMPA (80μM) and NMDA (200μM) have no significant effects on [Ca^2+^] in dendrons. Dotted lines indicate 95% confidence intervals. Numbers of dendrons (n) and mice (N) are given for each treatment and each sex.

Application of a 90s puff of kisspeptin (100 nM) generated a 35.1±0.1% (mean±SEM, male, median 35.1%) to 40.5±0.2% (female, median 40.1%) rise in dendron [Ca^2+^] that peaked and then gradually subsided across the 400 s duration of the recording in both male (N=7) and female (N=5) mice (Fig.2C,D). Approximately 91% of dendrons responded to kisspeptin.

In contrast, 90 s puffs of glutamate (600μM) that would activate both ionotropic and metabotropic receptors, were found to have no significant effects on dendron [Ca^2+^] in either males (N=9) or females (N=2)(Fig.2E,F). Furthermore, 90 s puffs of AMPA (80μM) had no effects on dendron [Ca^2+^] in males (N=5) or females (N=2)(Fig.2G,H). In further experiments (N=2 males, N=2-3 females), glutamate and AMPA were given as two 20 s puffs but were also found to have no effects on dendron fluorescence (not shown). Similarly, NMDA (200μM) given as two 30 s puffs in the absence of Mg^2+^ had no effects in males (N=4) or females (N=5)(Fig.2IJ).

We next tested the effects of the co-expressed neuropeptides NKB and dynorphin on the GnRH dendron. Ninety second puffs of 100 nM NKB generated small rises in dendron [Ca^2+^] (male, 2.45±0.04% (mean±SEM), median 2.54%; female, 4.73±0.05%, median 4.64%) that were not significantly different to control aCSF puffs in either males (N=11) and females (N=5)(Fig.3A,B). To ensure that this was not a technical false negative, horizontal brain slices were prepared from *Kiss1*-cre^+/-^;GCaMP6f mice in the exact same manner. Identical puffs of 100nM NKB were found to exert potent stimulatory effects on arcuate kisspeptin neuron [Ca^2+^] in both males (N=3) and females (N=3)(Fig.3C,D). The effects of NKB on arcuate kisspeptin neuron [Ca^2+^] were more potent than in males than females (area under curve = 18,542±2101 sec.%F versus 5,441±1,158, *P*<0.001, Mann-Whitney test)(Fig.3C,D).

**Figure 3.**
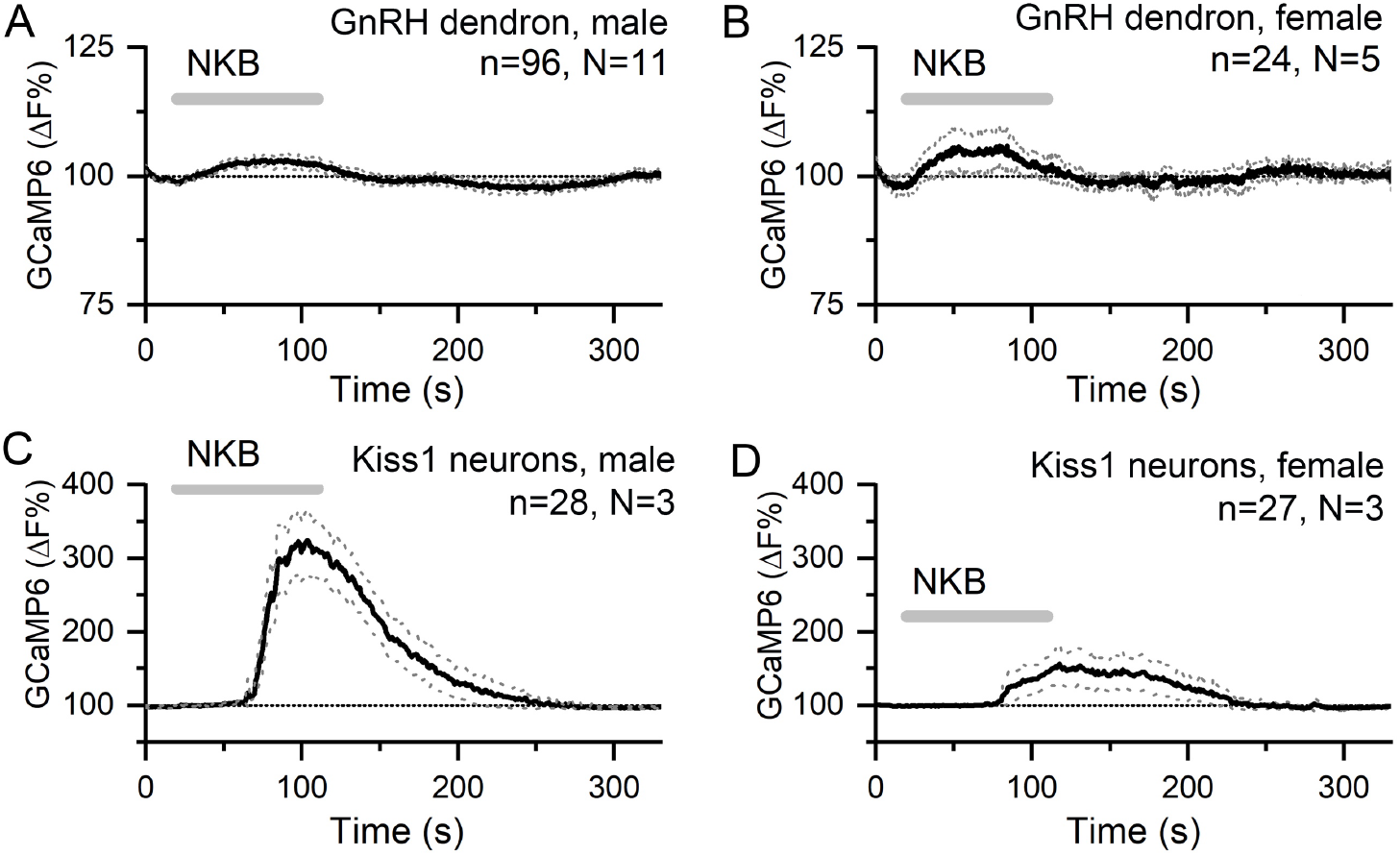
NKB increases [Ca^2+^] in KNDy neurons but not in GnRH neuron distal dendrons. **A&B**, 90s puffs of 100nM NKB have no significant effect on GCaMP6 fluorescence in GnRH neuron distal dendrons in male and female *Gnrh1-Cre::GCaMP6s* mice. **C&D**, 90s puffs of 100nM NKB evoke large, sexually differentiated increases in [Ca^2+^] in KNDy neurons of male and female *Kiss1-Cre;GCaMP6f* mice. Note the altered y-axis. Dotted lines indicate 95% confidence intervals. Numbers of dendrons (n) and mice (N) are given for each treatment and each sex.

Puffs of 200 nM dynorphin (90s) were also found to have no significant effect on dendron [Ca^2+^] in either males (N=11) and females (N=11)(male, 2.83±0.03%, median 2.86%; female, 2.14±0.05%, median 2.19%)(Fig.4A,B). As an inhibitory neuromodulator, it was possible that any suppressive actions of dynorphin may be difficult to assess on basal dendron [Ca^2+^]. As such, we tested the effect of dynorphin on kisspeptin-evoked increases in dendron [Ca^2+^]. However, dynorphin continued to fail to alter dendron [Ca^2+^] in males (N=5) and females (N=4)(Fig.4C,D).

**Figure 4.**
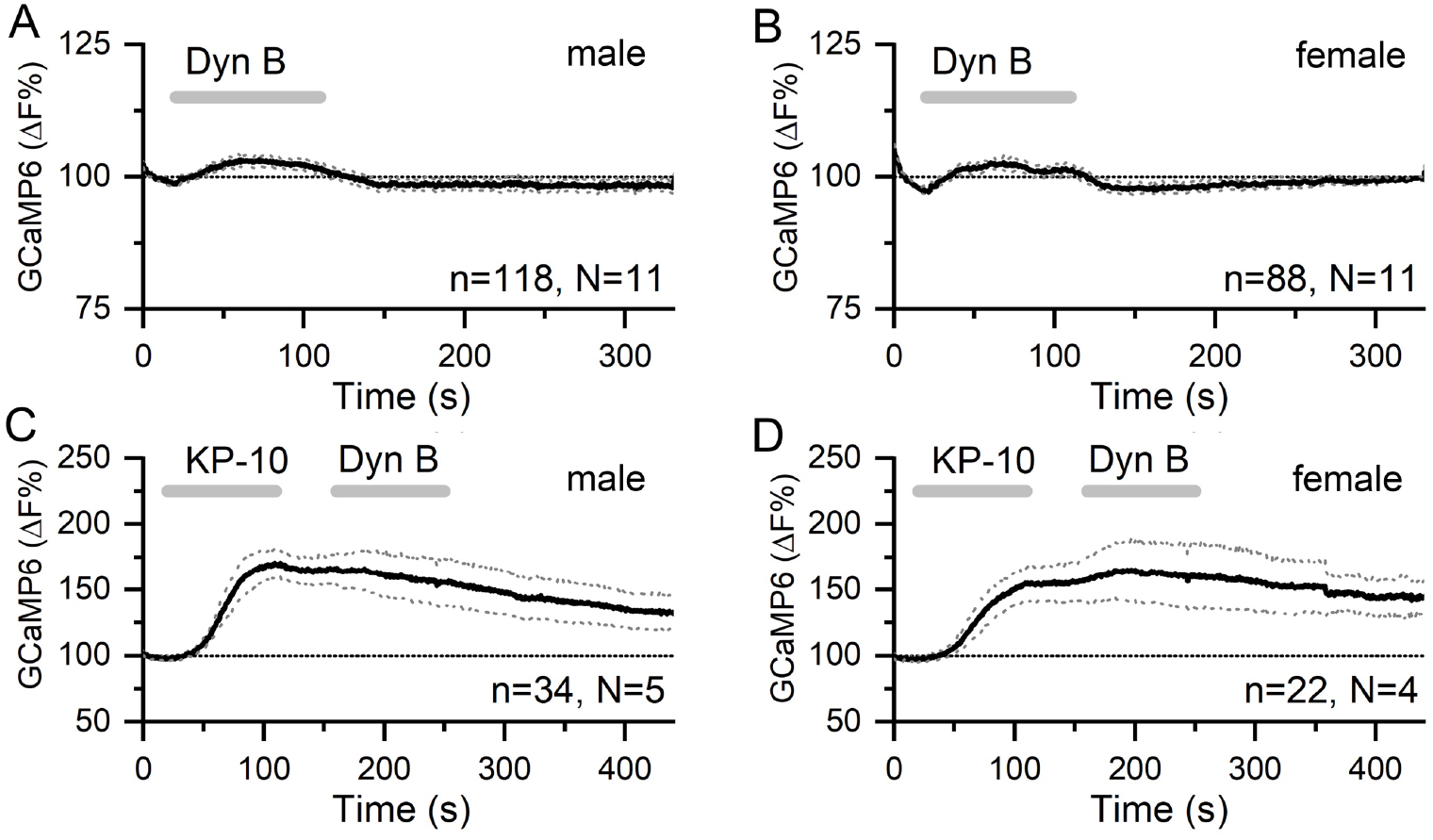
Dynorphin has no effect on [Ca^2+^] in GnRH neuron distal dendrons. **A&B**, 90s puffs of 200nM dynorphin have no significant effect on GCaMP6 fluorescence in GnRH neuron distal dendrons in male and female *Gnrh1-Cre::GCaMP6s* mice. **C&D**, similarly, dynorphin has no effect on kisspeptin-10-evoked increases in GCaMP6 fluorescence in either sex. Dotted lines indicate 95% confidence intervals. Numbers of dendrons (n) and mice (N) are given for each treatment and each sex.

### Kisspeptin signaling in KNDy neurons is not necessary for KNDy neuron synchronization

The experiments reported above indicate that, of the co-transmitters released from KNDy terminals, the GnRH neuron dendrons only express functional receptors for kisspeptin. This predicts that kisspeptin is the only co-transmitter released by KNDy terminals necessary to activate the GnRH neuron dendron and drive pulsatile LH secretion. This is in stark contrast to KNDy signalling at the KNDy cell body where evidence suggests the opposite relationship with kisspeptin having no role in synchronizing their activity while the co-released neuropeptides NKB and dynorphin form a recurrent collateral excitatory/inhibitory circuit critical for synchronization (de Croft, Boehm et al. 2013, Qiu, Nestor et al. 2016, Moore, Coolen et al. 2018).

To test this hypothesis *in vivo*, we generated mice in which only kisspeptin was deleted from KNDy neurons and assessed both KNDy neuron synchronization (transmission at the KNDy cell body) and pulsatile LH secretion (transmission at the KNDy terminals). In the homozygous state, the *Kiss1-Cre* mice used in this study represent a *Kiss-1* deletion (Yeo, Kyle et al. 2016). To characterize these mice further, immunohistochemical analyses of adult female *Kiss1*-Cre^- /-^ mice (N=4) confirmed the complete absence of kisspeptin peptide in cells or fibers within the ARN while immunoreactivity for NKB remained (Fig.5A-C).

**Figure 5.**
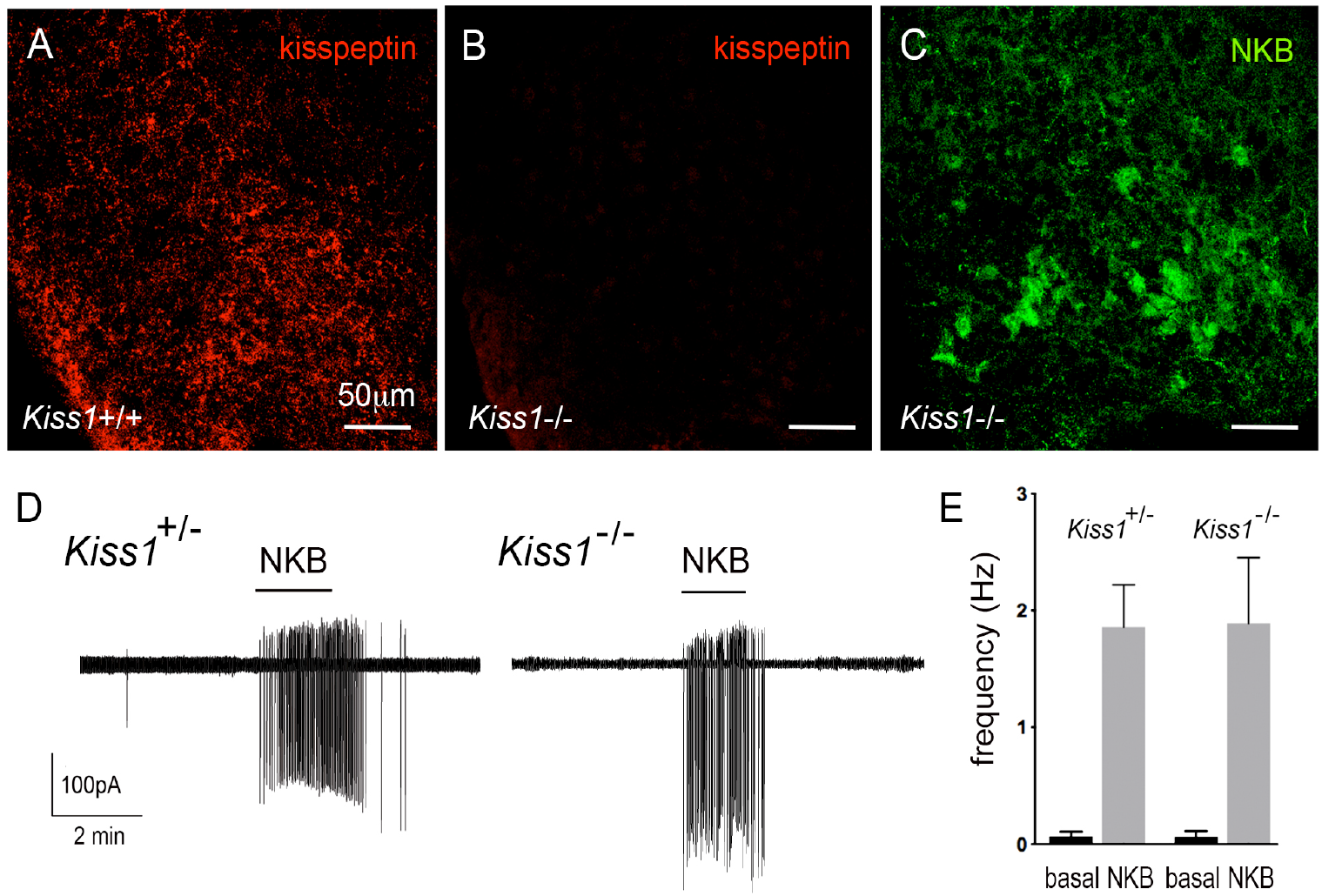
Characterization of Kiss1-null mice. **A-C**, immunofluorescence for kisspeptin (A,B) and NKB (C) in the ARN of wild-type (A) and *Kiss1-*null (B,C) female mice. **D**, Cell-attached recordings showing the effects of 100nM NKB on firing of KNDy neurons in acute brain slices prepared from female heterozygous and homozygous (null) *Kiss1-Cre;tdT* mice. **E**, mean±SEM changes in KNDy neuron firing evoked by 100nM NKB in heterozygous and homozygous (null) *Kiss1-Cre;tdT* mice.

We also assessed NKB receptor function at the KNDy neuron cell body in these mice by undertaking cell-attached recordings of KNDy neurons in the acute brain slice comparing the effects of 50 nM NKB on KNDy neuron firing in control *Kiss1-Cre*^*+/-*^;*tdT* (N=4) and null *Kiss1-Cre*^*+/-*^;*tdT* (N=5) mice. Both lines showed the same very low spontaneous firing rates (0.06±0.04Hz, n=14, *Kiss1-Cre*^*+/-*^, 0.06±0.06Hz, n=14, *Kiss1-Cre*^*-/-*^) typical of KNDy neurons (de Croft, Piet et al. 2012), and 50 nM NKB exerted the same marked stimulatory effect on firing (1.9±0.4Hz, n=14, *Kiss1-Cre*^*+/-*^, 1.9±0.6Hz, n=14, *Kiss1-Cre*^*-/*-^)(Fig.5D,E). Together, these studies indicate the normal expression of NKB and function of NKB receptors in KNDy neurons in the absence of kisspeptin in *Kiss1*-null mice.

The synchronized episodic activity of the ARN kisspeptin neurons can be measured in real time using *in vivo* GCaMP6 fiber photometry (Han, Clarkson et al. 2018). Twenty four hour GCaMP fiber photometry recordings from the middle/caudal ARN of adult female *Kiss1-Cre*^*-/-*^; *tdT* mice (N=4) given prior injections of AAV9-CAG-FLEX-GCaMP6s, revealed the presence of frequent abrupt ARN kisspeptin neuron synchronizations (Fig.6A). This pattern is similar to that observed in ovariectomized mice (McQuillan & Herbison, in preparation) and compatible with Kiss1-Cre^-/-^ mice being hypogonadal (Yeo, Kyle et al. 2016).

**Figure 6.**
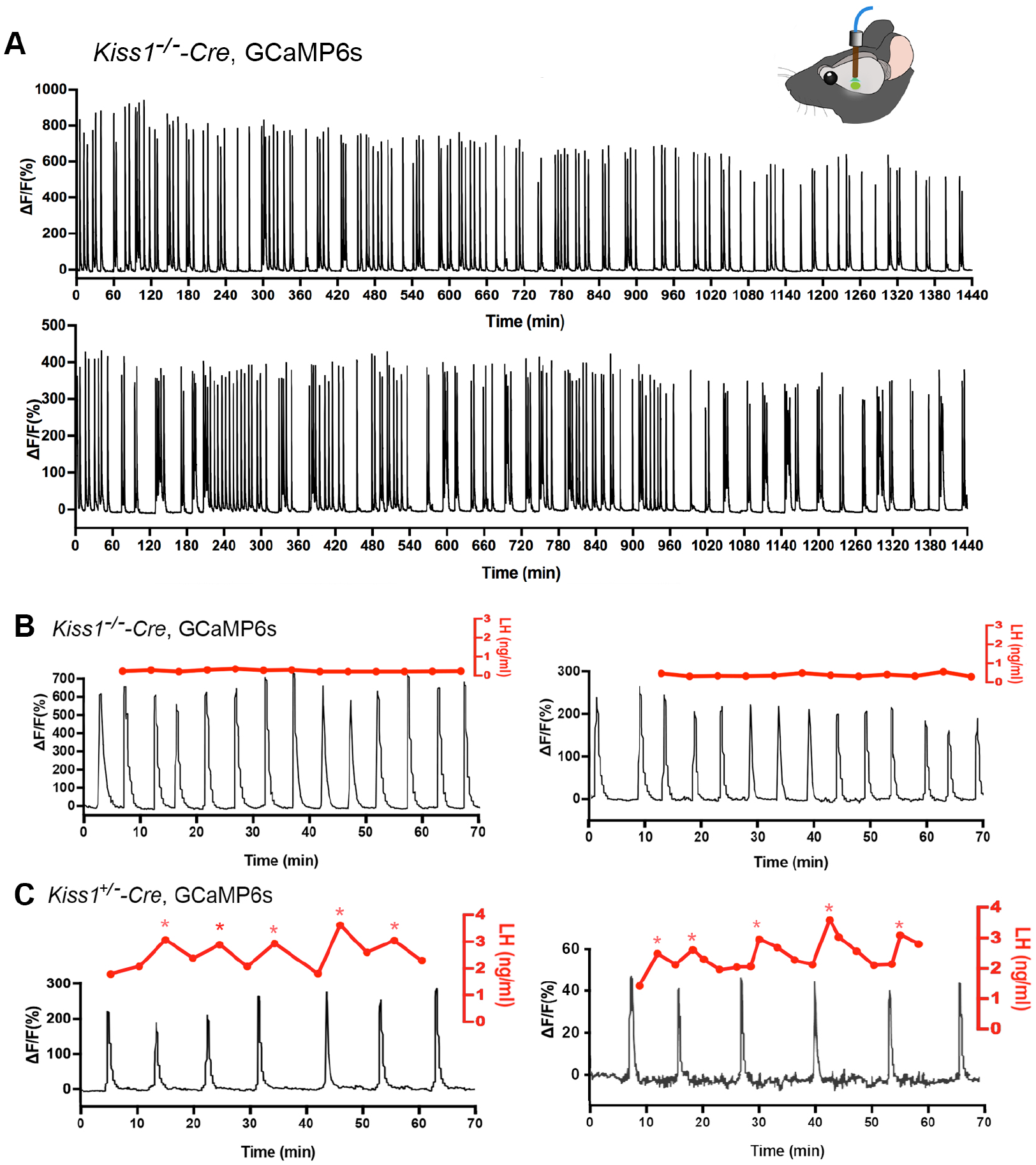
Mice with deleted Kiss1 exhibit KNDY neuron synchronization events but fail to generate pulsatile LH secretion. **A**, representative examples of 24h *in vivo* GCaMP6 fiber photometry recordings of KNDy neuron synchronization events from 2 female *Kiss1*^*-/-*^*-Cre;tdT::GCaMP6s* mice. **B**, representative examples of combined 5-minute tail-tip bleeding for LH levels (red) and GCaMP6 fiber photometry (black) recordings from 2 female *Kiss1*^*-/-*^*-Cre;tdT::GCaMP6s* mice. **C**, representative GCaMP6 photometry, 3-5 min tail-tip bleeding LH levels from 2 ovariectomized female *Kiss*^*+/-*^*-Cre;tdT::GCaMP6s* mice.

### Kisspeptin in KNDy neurons is essential for episodic activation of the GnRH neuron dendron to generate pulsatile LH secretion

The *in vitro* studies above indicate that kisspeptin is the only co-transmitter signaling from KNDy neurons to GnRH neurons. We reasoned that if this was the case then the synchronized output of KNDy neurons, that is normally perfectly correlated with pulsatile LH secretion (Han, Kane et al. 2019, McQuillan, Han et al. 2019), would fail to signal to the GnRH neuron dendron in *Kiss1*-null mice and result in an absence of pulsatile LH secretion. AAV-injected *Kiss1-Cre*^*-/-*^ mice (N=4) underwent fiber photometry recordings while also having 5-min tail tip bleedings performed for 1 hour to assess pulsatile LH secretion. As the nearest possible control, ovariectomized AAV-injected *Kiss1-Cre*^*+/-*^ mice (N=3) were assessed at the same time. Despite robust ARN^KISS^ neuron synchronization events, all four *Kiss1-*null mice exhibited an invariant, very low level of LH demonstrating a complete uncoupling of the KNDy pulse generator from LH secretion (Fig.6B). In contrast, all three ovariectomized *Kiss1-Cre*^*+/-*^ mice exhibited pulsatile LH secretion that was perfectly correlated with ARN kisspeptin neuron synchronization events (Fig.6C).

## DISCUSSION

Co-transmission in the brain is considered to result primarily from the differential release of synaptic vesicles containing separate neurotransmitters (van den Pol 2012, Vaaga, Borisovska et al. 2014, Tritsch, Granger et al. 2016). The KNDy neuron provides an example of co-transmission in which its output at recurrent collaterals and its primary efferent target is differentially interpreted by converse patterns of postsynaptic neuropeptide receptor expression (Fig.7). Kisspeptin is the only co-transmitter active at the GnRH neuron dendron whereas the exact opposite situation exists at KNDy neuron recurrent collaterals where all of the co-transmitters, except kisspeptin, are active (Navarro, Gottsch et al. 2011, de Croft, Boehm et al. 2013, Ruka, Burger et al. 2013, Qiu, Nestor et al. 2016). Thus, KNDy neurons conform to Dale’s Principle of uniform transmitter expression across their axonal arbor but solve the problem of differential signaling through opposite patterns of postsynaptic receptor expression at their targets (Fig.7). While this type of receptor-dependent co-transmission has been reported for small molecule transmitters it has not been shown for neuropeptides (Tritsch, Granger et al. 2016). We show that this mode of co-transmission is physiologically relevant *in vivo* as deletion of kisspeptin from the repertoire of transmitters used by KNDy neurons abolishes episodic hormone secretion while maintaining KNDy neuron synchronization behavior.

**Figure 7.**
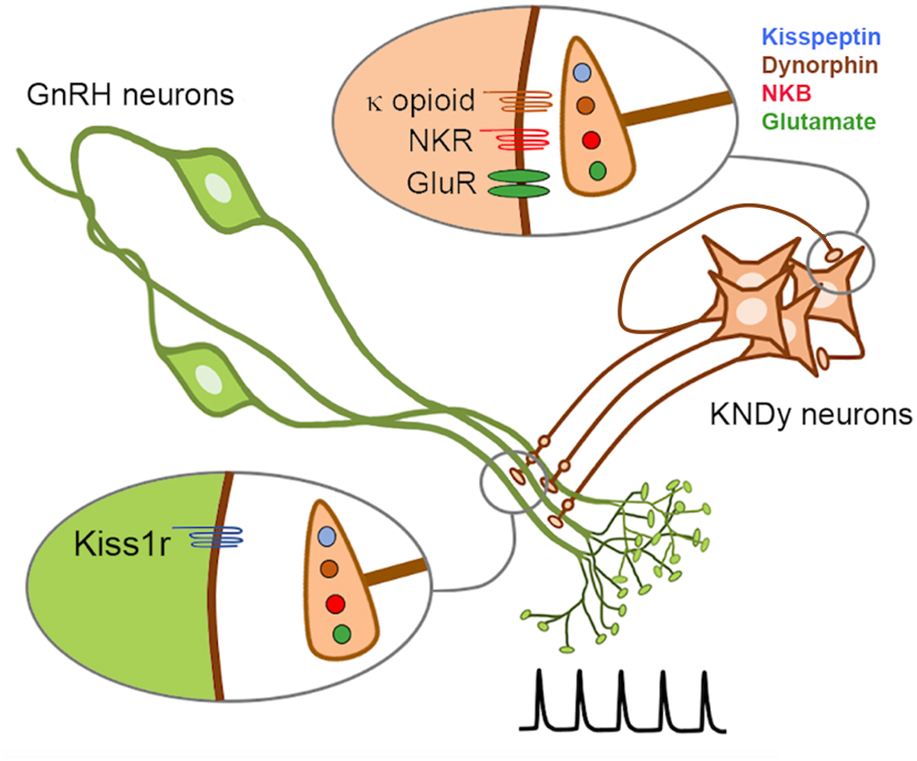
Schematic diagram depicting the proposed converse patterns of co-transmission that occur at the KNDy neuron recurrent collaterals and their projections to the GnRH neuron dendrons. NKR, neurokinin receptors; GluR, glutamate receptors.

As a result of their migration from the nose into the brain at mid-gestation, the GnRH neuron cell bodies are scattered throughout the basal forebrain (Wray 2010). The functional difficulties imposed by this topography appear to be solved, in part, by GnRH neurons focusing their blended dendritic/axonal projections on the median eminence where they can be regulated in a concerted manner by the KNDy pulse generator (Herbison 2018). While it is clear that classic synaptic inputs exist on the GnRH neuron dendron (Moore, Prescott et al. 2018, Wang, Guo et al. 2020), we note that this is not the case for the KNDy neuron innervation. Although many close appositions were identified with regular confocal analysis, no evidence was found with ExM for these to be synaptic inputs between KNDy fibers and GnRH neuron dendrons. This was despite the identification of *bona fide* kisspeptin synapses at the GnRH neuron cell body and proximal dendrites. An electron microscopic investigation in the rat has also reported that closely apposed kisspeptin fibers and GnRH neuron nerve terminals in the median eminence do not form synapses (Uenoyama, Inoue et al. 2011). This morphological relationship is indicative of short-distance volume transmission (van den Pol 2012). We also note that individual KNDy fibers form appositions with multiple GnRH neuron dendrons. Thus, it seems probable that the synchronous activation of many GnRH neuron dendrons is achieved by volume transmission from the multiple varicosities of KNDy neuron axons winding their way through GnRH neuron dendrons.

The nature of KNDy signaling at the GnRH neuron has been difficult to discern. Studies in the mouse show that KNDy neurons do not project to the GnRH neuron cell body and that the distal dendron is their only direct target (Yip, Boehm et al. 2015, Qiu, Nestor et al. 2016). Whereas immunohistochemical studies have indicated that GnRH neurons express receptors for NKB and/or dynorphin (Krajewski, Anderson et al. 2005, Weems, Witty et al. 2016), this has not been supported by mRNA profiling and electrophysiological studies (Mitchell, Prevot et al. 1997, Sannella and Petersen 1997, Navarro, Gottsch et al. 2011, Qiu, Nestor et al. 2016). We find here that NKB had no effects on [Ca^2+^] in GnRH neuron dendrons indicating that they are not likely to express TAC3R. This is in stark contrast with the KNDy neurons themselves where NKB evokes a strong activation in the same brain slice preparation. We also found no evidence for dynorphin to modulate either basal or kisspeptin-evoked [Ca^2+^] in the GnRH dendron. The latter result does not support the hypothesis that dynorphin signaling is involved in terminating the kisspeptin-evoked GnRH pulse at the level of the GnRH neuron, at least in mice (Weems, Coolen et al. 2018). Surprisingly, we observed that glutamate, AMPA and NMDA had no impact on [Ca^2+^] in GnRH neuron dendrons and it remains possible that small changes in [Ca^2+^] beyond those detectable with GCaMP6 exist. Nevertheless, functionally, it is notable that KNDy neurons need to be activated at >5Hz to achieve any significant episodic release of LH *in vivo* (Han, McLennan et al. 2015).

The KNDy neurons package their individual neurotransmitters into separate vesicles (Murakawa, Iwata et al. 2016) thereby allowing the possibility of differential trafficking of vesicles within the cell. However, this does not appear to occur as others (Lehman, Coolen et al. 2010) and ourselves have shown that the recurrent collaterals and projections to GnRH neuron dendrons all express kisspeptin, NKB and dynorphin. Further the optogenetic activation of KNDy neurons in brain slices has been demonstrated to release functionally relevant NKB and dynorphin (Qiu, Nestor et al. 2016). As such, it is interesting to consider whether the episodic release of NKB and dynorphin from KNDy varicosities in the region of the ventrolateral ARN may impact on other ARN neuronal cell types. Similarly, given that kisspeptin exerts both inhibitory and excitatory actions on other ARN neurons (Fu and van den Pol 2010, Liu and Herbison 2015), it is possible that abrupt episodes of kisspeptin release in the vicinity of the KNDy cell bodies may entrain other ARN networks to the GnRH pulse generator. The potential function or redundancy of these “off-network” signaling events remain to be established.

In summary, we provide evidence that the GnRH pulse generator network uses converse patterns of postsynaptic receptor expression at the two main targets of the KNDy neuron to drive episodic hormone secretion. This likely solves the problems of signal resolution generated when using neuropeptide volume transmission in the close vicinity of the KNDy somata and GnRH neuron dendrons. Although highly redundant at the level of the post-synaptic space, this novel pattern of co-transmission enables neurons to achieve differential signaling at varied targets without the need for selective trafficking of neuropeptide transmitters throughout their axonal arbor.

## Acknowledgements

Studies were supported by the New Zealand Health Research Council and Wellcome Trust. Dr. A. Caraty and Dr. P. Ciofi (University of Bordeaux) are thanked for generous gifts of antisera. The authors thank Dr. Karl Iremonger for commenting on an earlier version of the manuscript.

## Materials and Methods

### Animals

C57BL6 *GnRH-GFP* mice (Spergel, Kruth et al. 1999), C57BL6/J *Gnrh1-Cre* mice (JAX stock #021207)(Yoon, Enquist et al. 2005), 129S6Sv/Ev C57BL6 *Kiss1-Cre* mice (Yeo, Kyle et al. 2016) alone or crossed on to the Ai9-CAG-tdTom^+/-^ reporter line (JAX stock #07909)(Madisen, Zwingman et al. 2010)(*Kiss1-Cre;tdT* mice) or Ai95(RCL-GCaMP6f)-D line (JAX stock #028865)(Madisen, Garner et al. 2015) (*Kiss1-Cre;GCaMP6f* mice) were group-housed in individually-ventilated cages with environmental enrichment under conditions of controlled temperature (22±2°C) and lighting (12-hour light/12-hour dark cycle; lights on at 6:00h and off at 18:00h) with *ad libitum* access to food (Teklad Global 18% Protein Rodent Diet 2918, Envigo, Huntingdon, UK) and water. Daily vaginal cytology was used to monitor the estrous cycle stage. All animal experimental protocols were approved by the University of Otago, New Zealand or the University of Cambridge, UK.

### Stereotaxic surgery

Adult mice (2-4 months old) were anaesthetized with 2% isoflurane and placed in a stereotaxic apparatus with prior local Lidocaine (4mg/Kg bodyweight, s.c.) and Carprofen analgesia (5 mg/kg body weight, s.c.). A custom-made bilateral Hamilton syringe apparatus holding two syringes with needles held 0.9 mm apart was used to perform bilateral injections into the preoptic area (AP 0.10 mm, depth 4.3 mm) or ARN (AP −2.00 mm, depth 5.9 mm) of *Gnrh1-Cre* or *Kiss1-Cre;tdT* mice, respectively. The needles were lowered into place over 2 min and left *in situ* for 3 min before the injection was made. One μL of AAV9-CAG-FLEX-GCaMP6s-WPRE-SV40 (1.7 × 10-^13^ GC/mL, University of Pennsylvania Vector Core) was injected into the ARN of *Kiss1-Cre;tdT* mice and 1.5 μL into the preoptic area of *Gnrh1-Cre* mice at a rate of ∼100 nl/min with the needles left in situ for a further 10 min before being withdrawn. *Kiss1-Cre;tdT* mice were then implanted with a unilateral indwelling optical fiber (400 µm diameter; 0.48 NA, Doric Lenses, Quebec, Canada) positioned directly above the mid-caudal ARN using the same coordinates for the AAV injections. Carprofen (5 mg/kg body weight, s.c.) was administered for post-operative pain relief. Mice were housed individually for the remaining experimental period. A typical post-surgical resting period of 4-6 weeks was allowed, with daily handling and habituation.

### Immunohistochemistry for Expansion Microscopy

Diestrus *GnRH-GFP* mice aged between 2-4 months old were perfused transcardially with 4% PFA in 0.1 M phosphate-buffered saline. Fifty-micron coronal brain sections were cut on a vibratome and kept in cryoprotectant until used. Sections were pre-treated with 0.1% sodium borohydrate (Sigma-Aldrich) in Tris-buffered saline (TBS) for 15 min at RT and then further treated with 0.1% Triton-X-100 (Sigma-Aldrich) and 2% goat serum in TBS overnight at 4°C for improving antibody penetration. Next, sections were washed and incubated for 72 h at 4°C with a cocktail of chicken anti-GFP (1:8000; Abcam), guinea pig anti-synaptophysin1 (1:800; Synaptic Systems) and rabbit anti-kisspeptin-10 (1:2000; gift from Alain Caraty) antisera added to the incubation solution made up of TBS, 0.3% Triton-X-100, 0.25% bovine serum albumin and 2% goat serum. All subsequent incubations were performed using the same incubation solution. Sections were rinsed in TBS, followed by incubation with biotinylated goat anti-guinea pig immunoglobulin (Vector Laboratories) mixed with Alexa488-conjugated goat anti-chicken (Thermo Fisher Scientific) and ATTO 647N goat anti-rabbit immunoglobulins (Sigma-Aldrich), 1:400 each, for 15 h at 4°C. Sections were then expanded as previously reported (Wang *et al*., 2020 eLife). Briefly, sections were stained with 1:1000 Hoechst 33342 dye (Thermo Fisher Scientific), processed for linking with anchoring agent and trimmed to include the rostral preoptic area or arcuate nucleus region. Next, sections were incubated in monomer solution, followed by gelling solution in a humidified chamber at 37°C. Gel-embedded sections were digested overnight with Proteinase K. Following that, the gels were rinsed in PBS, incubated with 1:1500 Streptavidin-568 at 37°C for 3 h and rinsed again with PBS. Lastly, water was added every 20 min, up to 5 times during the expansion step. Expanded samples were placed in imaging chamber filled with water and cover slipped using #1.5 cover glass.

Imaging was undertaken using a Nikon A1R upright confocal microscope equipped with a water-immersing lens (25X Numerical Aperture 1.1; working distance 2 mm). All images were captured using sequential scanning mode using 500-550 nm, 580-620 nm and 620-660 nm bandpass filters for Alexa Fluor 488, Alexa Fluor 568 and ATTO 647, respectively. All image stacks (frames: 102.51 × 51.25µm, 1024 × 512 pixels) were acquired at 0.6 μm focus intervals. Images were analyzed using ImageJ to determine the number of kisspeptin terminals containing synaptophysin apposed to GnRH proximal or distal dendrites. For the cell body/proximal dendrite, 250μm (60μm pre-expansion) of contiguous primary dendrite arising from the GnRH cell body was randomly selected from three rostral preoptic area sections in each of the 3 mice. Each apposing synaptophysin-kisspeptin bouton (diameter >0.4μm) was examined to establish the side-on view or a z-stack/face-view of the imaged synapse. A line scan was then performed across this plane and the relative intensity of the Alexa488 and ATTO647 was measured and plotted in Microsoft Excel. For the face-view orientation, a 2 µm x 2 µm (width x length) box was drawn on each of the synaptophysin-kisspeptin boutons to measure the relative intensity of Alexa488 and ATTO647 in z-stack through scan. An apposition was considered a synapse where the signals overlapped by >0.95 μm (0.23 μm pre-expansion) in the side-on plane or >1.75 μm (0.42 μm pre-expansion) in the z-stack through/face-view (Wang, Guo et al. 2020). The same method was used to establish the relationship between synaptophysin-kisspeptin profiles and the distal dendron using 60 μm (15 μm pre-expansion) contiguous lengths of dendron located in the ventrolateral ARN.

### Immunohistochemistry for multi-label fluorescence

Adult diestrus *Kiss-Cre*^*+/-*^ and *Kiss1*-Cre^-/-^ mice (2-3 months old) underwent transcardial perfusion. Coronal brain sections of 40 μm thickness were prepared and incubated in rabbit anti-NKB (1:5000; Novus Biologicals) and sheep anti-Kisspeptin 10 antisera (1:8000; gift of Alain Caraty) followed by biotinylated donkey anti-sheep immunoglobulins (1:1500; Jackson Immunoresearch), donkey anti-rabbit conjugated with Alexa Fluor 488 (1:1000, Thermo Fisher Scientific) and Streptavidin Alexa Fluor 647 (1:1500, Thermo Fisher Scientific). Images were acquired using a Leica SP8 Laser Scanning Confocal Microscope and a 63x oil immersion objective (Numerical Aperture 1.20; Working Distance 300μm) with image stacks collected at 0.6μm intervals (Cambridge Advanced Imaging Centre).

Adult *GnRH-GFP* mice were perfused transcardially and a ventral para-horizontal brain slice prepared by making a single horizontal cut from the base of the brain (Suppl.Fig.1). Brain slices were incubated in rabbit anti-kisspeptin 10 (1:2000; gift of Dr. Alain Caraty), guinea pig anti-NKB (1:5000; IS-3/61, gift of Dr. Philippe Ciofi) and chicken anti-GFP (1:2000; AB16901, Chemicon) antisera followed by goat anti-chicken Alexa Fluor 488, (1:200, A-11039, Thermo Fisher) goat anti-guinea pig Alexa Fluor 568 (1:200, A-11075, Invitrogen) and goat anti-rabbit Alexa Fluor 633, (1:200, A-21071, Invitrogen) immunoglobulins. Sections were examined on a Zeiss LSM 710 confocal microscope with a 63x/1.4 Plan Apochromat objective at 1.4x zoom and Nyquist resolution using ZEN software (version 5,5,0,375). Image stacks were acquired at 0.38 µm intervals. The ventrolateral ARN was imaged at 0.38 µm z-step intervals with 4 image stacks tiled 2×2 taken at 1056×1056 pixels resolution and stitched using the 3D Stitching plugin in FIJI in linear blend mode (Preibisch, Saalfeld et al. 2009). For each animal, 10 regions of >100 µm length of GFP-labeled dendrons selected due to their proximity to the ME and direction of projection towards ME were analyzed. A close apposition was defined as the absence of dark pixels between elements or even a slight overlap of the different channels. For 3-dimensional reconstruction, raw data image stacks were isosurface rendered in Amira (version 5.3, Visage Imaging, San Diego) using the neuronal reconstruction plugin by (Schmitt, Evers et al. 2004).

### Confocal imaging of GnRH neuron dendron Ca^2+^ concentration

The procedure for making GCaMP6 recordings from GnRH neuron distal dendrons has been reported previously (Iremonger, Porteous et al. 2017). In brief, the mouse was killed by cervical dislocation, the brain quickly removed and optic tract peeled off. The dorsal surface of the brain was then glued to a vibratome cutting stage (VT1200s, Leica) and submerged in ice-cold (<2°C) sucrose-containing cutting solution (75mM NaCl, 75 sucrose, 2.5 KCl, 20 HEPES, 15 NaHCO_3_, 0.25 CaCl_2_, 6 MgCl_2_, 25 D-glucose, bubbled with 95% O_2_/5% CO_2_; 320 mOsmol). A single 500-µm-thick horizontal slice containing the median eminence and surrounding tissue (Suppl. Fig.1) was made and incubated in the cutting solution (34±1°C) and then recording aCSF (118mM NaCl, 3 KCl, 10 HEPES, 25 NaHCO_3_, 2.5 CaCl_2_, 1.2 MgCl_2_, 11 D-glucose; 95% O_2_/5% CO2; 27±1°C) for at least 1h before being transferred to the recording chamber (27±1°C) where the slice was held between two meshes with a perfusion flow rate of 1.5 ml/min. Imaging was performed with an Olympus FV1000 confocal microscope fitted with a 40X, 0.8 NA objective lens and 3X zoom with the aperture fully open. GCaMP was excited with a 488 nm Argon laser and emitted light passed through a 505–605 nm bandpass filter.

Test compounds were dissolved in aCSF and locally puff-applied with a patch pipette (4–6 MΩ) at low pressure (∼1 psi) controlled by a Pneumatic Picopump (PV821, World Precision Instruments, USA) for 20 to 90s. Total recording time for each test was 500s. The tip of the puff pipette was positioned 30 to 130µm above the surface of the slice. Initially, GnRH neuron dendrons were tested with 30s puffs of 20mM KCl to assess their viability and only those fibers displaying fast Ca^2+^ responses were used for experiments. Puffs of aCSF alone could generate small changes in signal with a slow rise and decay that followed the timing of the puff (4.50±0.05%, range of −12 to 12% with a median of 4.44%). Acceptable recordings had to have <5% drift in baseline fluorescence intensity across the time of the experiment. Control aCSF recordings were undertaken throughout the series of experiments and kisspeptin tests were undertaken as the last challenge. To block action potential-dependent synaptic transmission, TTX (0.5 to 1μM; Alamone Labs, Israel) was added to the recording aCSF for the duration of the experiment. For neuropeptide tests, GABAzine (5μM; Tocris Bioscience, UK), D-AP5 (50μM, Tocris) or DL-AP5 (25μM) and CNQX (10μM, Tocris) were added to the TTX-containing aCSF to eliminate ionotropic GABA_A_ receptor and glutamate receptor activation. A 0mM Mg^2+^ aCSF containing TTX was applied at least 5min before NMDA (200 μM; Tocris) puffs. Glutamate (600 μM; Tocris), AMPA (80μM; Tocris), Neurokinin B (100 nM; Tocris), Dynorphin B (100-200 nM; Tocris) and Kisspeptin (100 nM, Calbiochem, USA) were maintained as frozen stock solutions and dissolved into the recording aCSF.

Image acquisition was performed with Fluoview 1000 software. Frame scans (512×512 pixels) were performed on zoomed regions at 0.9Hz frame rate with the lowest possible laser power. Image analysis was performed with Fluoview1000 software and ImageJ. Regions of interest (ROI) were drawn around individual GnRH neuron dendrons. The percentage of GCaMP6 fluorescence was calculated as GCaMP6 ΔF%= 100x(F/F0), where F is the average fluorescence of ROI in each consecutive frame and F0 is the average fluorescence of ROI in first 10-20 frames before test drug puffs. Any tissue drift in the x-y axis was corrected by enlarging the ROI or by using Turboreg (ImageJ). All data are presented as mean±SEM. The average calcium traces are shown with lower and upper 95% coefficients and have a minimum animal number of 4. Statistical analyses were performed with non-parametric Paired sample Wilcoxon Signed Ranks Test or repeated-measures tests (Kruskal–Wallis or Friedman test with a post-hoc Dunn’s Test).

### Acute brain slice electrophysiology

Brain slice electrophysiology was undertaken as reported previously (Hessler, Liu et al. 2020). In brief, adult male and diestrous-stage female *Kiss1-Cre;tdT* mice were killed by cervical dislocation and 250µm-thick coronal brain slices containing the ARN prepared on a vibratome (VT1000S; Leica) in an ice-cold 75mM sucrose aCSF cutting solution. Slices were then incubated for at least 1h in recording aCSF (120mM NaCl, 3 KCl, 26 NaHCO_3_, 1 NaH_2_PO_4_, 2.5 CaCl_2_, 1.2 MgCl_2_, 10 HEPES, 11 glucose 95%O_2_/5%CO2, 32°C) before being transferred to a recording chamber. Loose-seal cell-attached recordings (10–30 MΩ) were made from tomato-expressing KNDy neurons visualized through an upright BX51 Olympus microscope. Cells were visualized by brief fluorescence illumination and approached using infrared differential interference contrast optics. Loose seals were made using recording electrodes (3.5-5.2 MΩ) filled with aCSF and action currents recorded in the voltage clamp mode with a 0mV voltage command. Signals were recorded using a Multiclamp 700B amplifier (Molecular Devices, Sunnyvale, CA) connected to a Digidata 1440A digitizer (Molecular Devices) and low-pass filtered at 3 kHz before being digitized at a rate of 10 kHz. For analysis, spikes were detected using the threshold crossing method. Signal acquisition and analysis was carried out with pClamp 10.7 (Molecular Devices). The effects of NKB on KNDy neurons were assessed by adding 50nM NKB to the aCSF for 1 min.

### In vivo GCaMP photometry and LH pulse bleeding

The GCaMP6 fiber photometry procedure used to detect KNDy neuron synchronization events (SEs) has been described in full previously (Clarkson, Han et al. 2017, Han, Kane et al. 2019, McQuillan, Han et al. 2019). In brief, ∼69% of KNDy neurons express GCaMP6 with this AAV approach and 96% of all GCaMP-expressing cells in the ARN are kisspeptin neurons. Freely-behaving AAV-injected *Kiss1*^*-/-*^*-Cre;tdT* mice were connected to the fiber photometry system for 4-6h using a fiber optic patch cord with fluorescence signals measured using a scheduled mode (5s on, 15s off). SEs were defined as peaks in fluorescence greater than 10% of maximum signal, thereby accounting for inter-animal variation in signal strength. Processed fluorescence signals were calculated as ΔF/F (%), = 100 x (Fluorescence - basal Fluorescence)/Fluorescence. To examine the relationship between SEs and pulsatile LH secretion, ∼1.5h of photometry recording was coupled with 5-min interval tail-tip blood sampling (4μL) as reported previously (Clarkson, Han et al. 2017, Han, Kane et al. 2019, McQuillan, Han et al. 2019). Levels of LH were measured by ELISA with an assay sensitivity of 0.04 ng/mL and intra-assay coefficient of variation of 9.3%.

## Key Resources Table

| Key Resources Table |  |  |  |  |
| --- | --- | --- | --- | --- |
| Reagent type (species) or resource | Designation | Source or reference | Identifiers | Additional information |
| Genetic reagent (*M. musculus*) | STOCK Tg(Gnrh1-cre)1Dlc/J | Jackson Laboratory | Stock #: 021207<br>RRID:IMSR_JAX:021207 |  |
| Genetic reagent (*M. musculus*) | B6.DBA-Tg(Gnrh1-EGFP)1Phs | Spergel et al., doi: 10.1523/JNEUROSCI.19-06-02037.1999 | MGI:6158458 |  |
| Genetic reagent (*M. musculus*) | Kiss1<sup>tm2(CreGFP)Coll.</sup>:tdTom | Yeo et al., doi:org/10.1111/jne.12435 2016 |  |  |
| Genetic reagent (*M. musculus*) | B6.Cg-Gt(ROSA)26Sor<sup>tm9(CAG-tdTomato)Hze</sup>/J (Ai9) | Jackson Laboratory | Stock #: 07909<br>RRID:IMSR_JAX:007909 |  |
| Genetic reagent (*M. musculus*) | B6J.Cg-Gt(ROSA)26Sor<sup>tm95.1(CAG-GCaMP6f)Hze</sup>/MwarJ (Ai95D) | Jackson Laboratory | Stock #: 028865<br>RRID:IMSR_JAX:028865 |  |
| transfected construct (*M. musculus*) | AAV9-CAG-FLEX-GCaMP6s-WPRE-SV40 | Penn Vector Core | RRID:Addgene_100844 | 1.7 x 10<sup>-13</sup> GC/mL |
| Antibody | Anti-Kisspeptin 10 (rabbit polyclonal) | Alain Caraty, INRA, France | Cat# AC566<br>RRID: AB_2314709 | (1:2000) |
| Antibody | Anti-Kisspeptin 10 (sheep polyclonal) | Alain Caraty, INRA, France<br>Franceschini *et al.*, 2013 | Cat# AC024 | (1:8000) |
| Antibody | Anti-NKB (rabbit polyclonal) | Novus Biologicals | Cat# NB300-201<br>RRID: AB_10000783 | (1:5000) |
| Antibody | Anti-NKB (guinea pig polyclonal) | P. Cioffi; INSERM; France | Cat# IS-3/63<br>RRID: AB_2732894 | (1:5000) |
| Antibody | Anti-GFP (chicken polyclonal) | Abcam | Cat# AB13970<br>RRID: AB_300798 | (1:8000) |
| Antibody | Anti-Synaptophysin 1 (guinea pig polyclonal) | Synaptic Systems | Cat#101004<br>RRID: AB_1210382 | (1:800) |
| Antibody | Goat anti-chicken<br>(goat polyclonal,<br>Alexa488-conjugate) | Thermo Fisher<br>Scientific | Cat# A-11039<br>RRID: AB_2534096 | (1:400) |
| Antibody | Goat anti-guinea pig<br>(goat polyclonal,<br>biotin-conjugated) | Vector<br>Laboratories | Cat# BA-7000<br>RRID: AB_2336132 | (1:400) |
| Antibody | Goat anti-rabbit<br>(goat polyclonal,<br>ATTO647N-conjugated) | Sigma-Aldrich | Cat# 40839<br>RRID: AB_1137669 | (1:400) |
| Antibody | AffiniPure F(ab')<sub>2</sub><br>Fragment Donkey<br>Anti-Sheep (donkey<br>polyclonal, biotin-conjugated) | Jackson<br>ImmunoResearch<br>Labs | Cat# 713-066-147<br>RRID: AB_2340717 | (1:1500) |
| Antibody | Alexa Fluor 488-F(ab')<sub>2</sub> Fragment<br>Donkey Anti-Rabbit<br>(H+L) | Jackson<br>ImmunoResearch<br>Labs | Cat# 711-546-152<br>RRID: AB_2340619 | (1:1000) |
| Antibody | Alexa Fluor 488 Goat<br>anti-chicken IgY (H+L) | Thermo Fisher | Cat# A-11039<br>RRID: AB_2534096 | (1:200) |
| Antibody | Alexa Fluor 568 goat<br>anti-guinea pig IgG<br>(H+L) | Invitrogen | Cat# A-11075<br>RRID: AB_2534119 | (1:200) |
| Antibody | Alexa Fluor 633 goat<br>anti-rabbit IgG (H+L) | Invitrogen | Cat# A-21071<br>RRID: AB_2535732 | (1:200) |
| chemical<br>compound, drug | (RS)-α-amino-3-hydroxy-5-methyl-4-isoxazolepropionic<br>acid (AMPA)) | Tocris | Cat#: 0169 | 80μM |
| chemical<br>compound, drug | D-2-Amino-5phosphonopentanoi<br>c acid sodium salt (D-AP5) | Tocris | Cat#: 0106/1 | 50μM |
| chemical<br>compound, drug | DL-2-Amino-5phosphonopentanoi<br>c acid sodium salt<br>(DL-AP5) | Tocris | Cat#: 3693/10 | 25μM |
| chemical<br>compound, drug | 6-Cyano-7nitroquinoxaline-2,3dione disodium<br>(CNQX) | Tocris | Cat#: 1045/1 | 10μM |
| chemical<br>compound, drug | Dynorphin B (Dyn B) | Tocris | Cat#:3196/1 | 100-200nM |
| chemical compound, drug | SR 95531 hydrobromide (GABAazine) | Tocris | Cat#: 1262/10 | 5µM |
| chemical compound, drug | Glutamate acid | Sigma-Aldrich | Cat#: G1251 | 300 or 600µM |
| chemical compound, drug | Kisspeptin-10 (KP-10) | Calbiochem | Cat#:45888 | 100nM |
| chemical compound, drug | Neurokinin B (NKB) | Tocris | Cat#:1582/1 | 50-100nM |
| chemical compound, drug | N-Methyl-D-aspartic acid (NMDA) | Tocris | Cat#: 0114 | 100 or 100µM |
| software, algorithm | Tetrodotoxin (TTX) | Alomone Labs | Cat#: T-550 | 0.5-2µM |
| software, algorithm | ImageJ image analysis software | ImageJ (https://imagej.net/) | RRID:SCR_003070 |  |
| software, algorithm | Vaa3D data visualization software | Vaa3D (http://www.vaa3d.org) | RRID:SCR_002609 |  |

---

**Suppl. Fig. 1.**
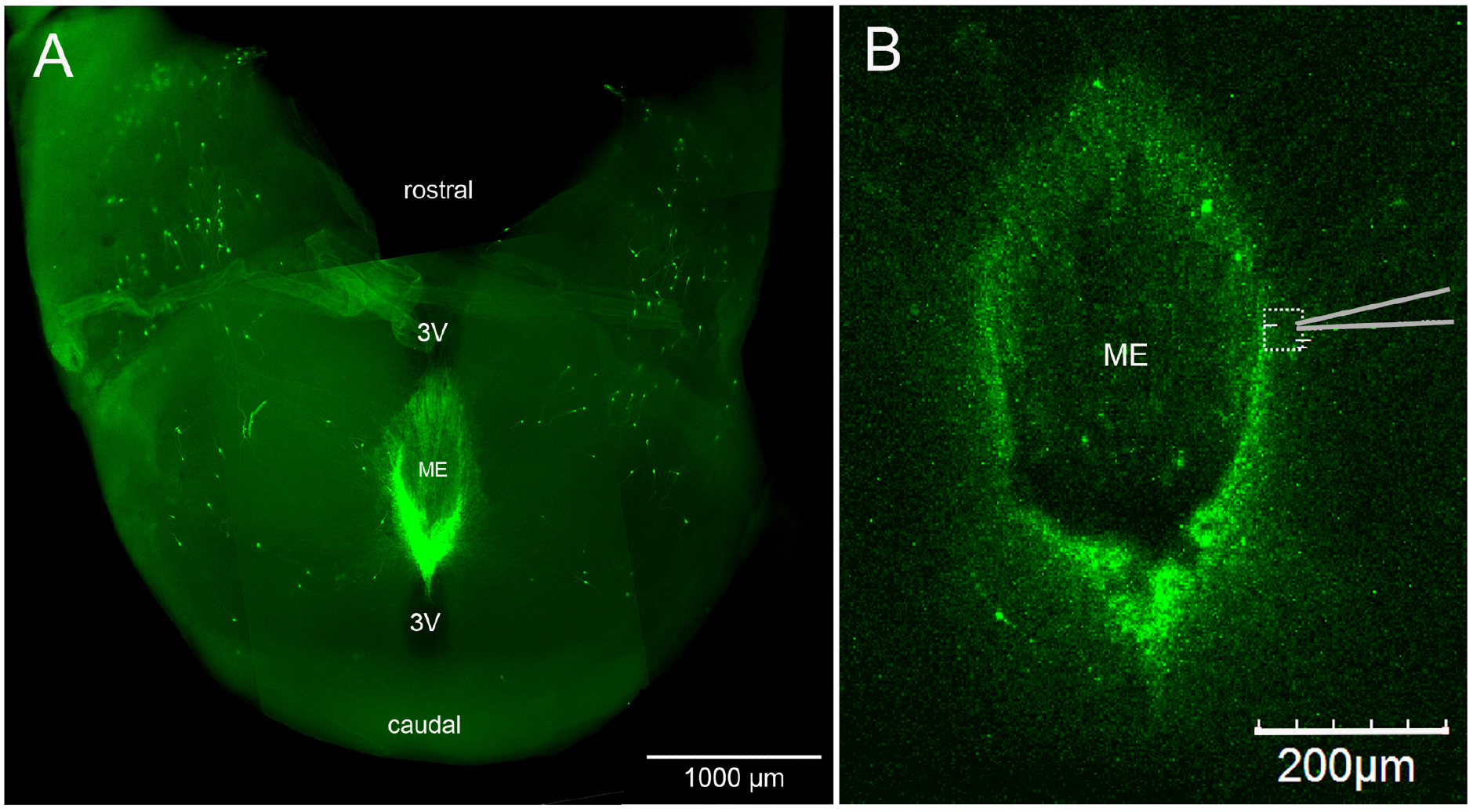
**A**. Looking down on a thick horizontal hypothalamic brain slice from a GnRH-GFP mouse in which the laterally positioned GnRH neurons of the anterior hypothalmaus and concentrated GnRH projections to the median eminence (ME) are visible. **B**. Higher-power view of the same orientation in a living slice from a GCaMP6s AAV-injected GnRH-Cre mouse showing the recording location (dotted square) and position of the puff pipette. 3V, third ventricle.

